# Biologically Inspired Deep Neural Network Models for Visual Emotion Processing

**DOI:** 10.1101/2025.10.20.683439

**Authors:** Peng Liu, Ke Bo, Yujun Chen, Andreas Keil, Mingzhou Ding, Ruogu Fang

**Author notes:** (MD) (RF).

## Abstract

The perception of opportunities and threats in complex visual scenes represents one of the main functions of the human visual system. The underlying neurophysiology is often studied by having observers view pictures varying in affective content. While deep neural networks (DNNs) have shown promise in modeling visual recognition of objects, their capacity to model visual affective processing remains to be better understood. In this study, we proposed a biologically inspired deep neural network model, referred to as the Visual Cortex Amygdala (VCA) model, that integrates a vision transformer with an amygdala-mimetic module designed to reflect both the anatomical hierarchy and self-attention-based computational mechanisms of the human brain. We evaluated the model along three dimensions: (1) predictive accuracy for emotional valence and arousal, (2) representational alignment with human amygdala activity, and (3)neural mechanisms of emotion representation. Our results showed that (1) the model can predict with high accuracy human emotion ratings on 1,182 images from the International Affective Picture System (IAPS) dataset (valence: *r* ≈ 0.9; arousal: *r* ≈ 0.7), (2) the model’s internal representations aligned with functional Magnetic Resonance Imaging (fMRI) data from the human amygdala, and (3) at the single neuron level, the amygdala module evolved emotion selectivity, and at the neuronal population level, the representational geometry became progressively more aligned with psychological models of emotion. Additional issues explored included (1) the effect of visual encoding and (2) the effect of structure and computational mechanisms on emotional assessment.

## Introduction

Visual experiences are not merely seen—they are also felt. A tranquil landscape, a threatening face, or a nostalgic object can evoke immediate, visceral affective responses. These associations between visual input and the emotional state it evokes constitute a foundational component of human cognition (Pessoa 2008; Barrett and Bar 2009). Yet, the neural mechanisms through which the emotional meaning of visual input is recognized remain incompletely understood, particularly concerning the interplay among perceptual processing, computational mechanisms, and anatomical constraints (Zald 2003; Vuilleumier 2005; Vuilleumier and Pourtois 2007; Kragel and LaBar 2016; Janak and Tye 2015).

Recent advances in artificial neural networks have provided a promising framework for exploring these questions. Deep neural networks (DNNs), including convolutional and transformer-based architectures, have achieved impressive success in modeling visual cognition. In particular, such models, when trained on large-scale affective image or multimodal datasets, can predict human emotional responses to visual stimuli. For instance, EmoNet (Kragel et al. 2019) formalized a mapping between visual features and emotional categories, and the more recent work by (Conwell et al. 2025) demonstrated that various DNNs, especially for CLIP-pretrained vision recognition models, could explain a substantial portion of the variability in human valence and arousal ratings (Radford et al. 2021). We recently showed that emotionally selective neurons can emerge even in deep neural networks trained solely on visual object recognition (Liu et al. 2024). These findings demonstrate the potential of using deep neural networks to model visual emotion processing. Further progress depends on a more systematic understanding of how architectural structure, training data modality, and computational mechanisms influence model performance.

Affective neuroscience has long recognized the amygdala as a central hub for emotion processing (LeDoux 2000; Phelps and LeDoux 2005). Its anatomical organization into hierarchically arranged subnuclei supports a broad range of affective functions, including rapid sensory appraisal, associative learning, and adaptive behavioral control (Swanson and Petrovich 1998; Janak and Tye 2015). Within this architecture, diverse cellular constituents, including excitatory and inhibitory neurons, as well as glial cells such as astrocytes, collaboratively shape the encoding and modulation of emotional information (Mederos and Perea 2019; Pacholko et al. 2020; Kaul et al. 2022; Seo et al. 2024). Emerging research suggests that astrocytes, traditionally regarded as passive support cells, may play an active computational role in information processing (Khakh and Sofroniew 2015; Yu et al. 2018). Through mechanisms such as tripartite synapse dynamics and slow calcium signaling, astrocytes are hypothesized to implement context-sensitive computations akin to self-attention operations, a key construct in the Transformer architecture, that influence circuit-level computation (Min et al. 2012; De Pitta’ et al. 2012; Vaswani et al. 2017; Gong et al. 2024). The existing deep learning models of emotion inference have not taken into account the amygdala’s hierarchical structure, cellular diversity, and emergent neural mechanisms. Embedding both anatomical and algorithmic principles into neural network models of the amygdala may enable more robust and biologically interpretable models of visual emotion perception.

In this study, we sought to develop and test a deep neural network model for visual emotion inference. The Visual Cortex Amygdala (VCA) model has two connected modules. The visual module consists of a vision transformer pretrained on image-text pairs (CLIP-ViT) and an amygdala-mimetic module incorporating known amygdala anatomy and self-attention-based computations (Wu et al. 2024; Di Chiano et al. 2025). Trained on multiple affective image datasets, including NAPS etc. and tested on 1,182 images from the International Affective Picture System (IAPS), the model achieves high predictive accuracy for emotional valence and arousal and maps accurately their joint distribution in the two-dimensional space spanned by valence and arousal (Bradley and Lang 2007). Importantly, through learning, the amygdala-mimetic module develops internal representations that align with human amygdala fMRI, and its units develop emotion selectivity consistent with the dimensional account of emotion (Posner et al. 2005; Rubin and and Talarico 2009; Putnam and Gothard 2019; Gothard 2020; Stanisławski et al. 2021). By comparing the performance of using different visual and amygdala modules in the VCA model we further examined the impact of visual encoding and amygdala architecture/computational mechanisms on emotional inference.

## Results

### The VCA model

The proposed VCA model consists of a visual module and an amygdala module, connected in a feed-forward fashion going from the visual cortex to the amygdala, a key route in visual emotion perception (Fig. 1a). The visual system module is a deep neural network pretrained on millions of text-image pairs, which encodes rich visual features and semantic information. The parameters in this module were held frozen in all experiments. The amygdala module, drawing inspiration from the structure and function of the biological amygdala, contains transformer self-attention blocks that represent distinct amygdala nuclei clusters, including the Lateral Nucleus (LA), Accessory Basal Nucleus (AB), Basolateral Nucleus (B), and Central Amygdala (CeA). In addition to the biologically inspired architecture, the module also incorporates an astrocyte-like computing mechanism within its self-attention layers, which is motivated by astrocyte-neuron interactions and dynamically weighs different input features in a way analogous how the brain focuses on various aspects of an emotional stimulus. We trained the amygdala module and its connections with the visual module in a supervised fashion to simultaneously predict valence (positive or negative emotion) and arousal (emotional intensity).

**Fig. 1.**
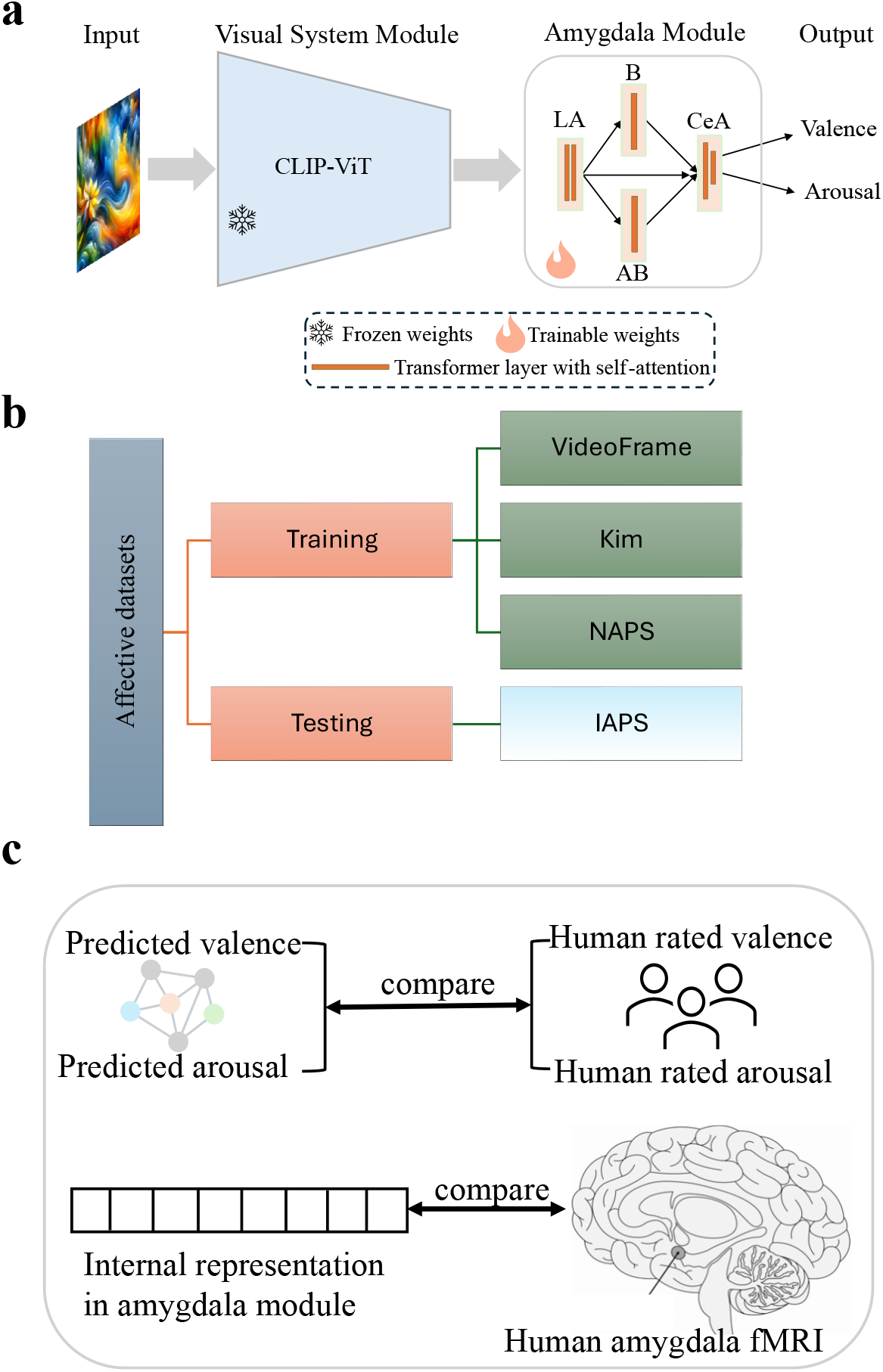
Study overview. **a**. Overview of the VCA model structure. An affective image is input into a visual system module consisting of a pre-trained neural network with frozen weights. The visual system output is then processed by the amygdala module, which includes the lateral nucleus (LA), accessory basal nucleus (AB), basolateral nucleus (B), and central amygdala (CeA) layers. The amygdala module’s output is the image’s valence and arousal. The structure of the amygdala module is inspired by the gross anatomy of the amygdala, while its computational mechanisms draw from astrocyte-like self-attention processes in the amygdala. **b**. Training and testing datasets used in this study. **c**. Model evaluation. The network’s predicted valence and arousal are compared to the normative valence and arousal ratings by human observers, and the internal representations in the amygdala module are compared to fMRI activity in the human amygdala.

The VCA model was trained using emotional images from three datasets: VideoFrame, Kim, and NAPS (Marchewka et al. 2014; Cowen and Keltner 2017; Kim et al. 2018) and tested the IAPS dataset (Fig. 1b). By training and testing the model on different datasets we sought to stress the generalizability of the emotion recognition capability of the model. In addition, the internal representations in the amygdala module were compared with human amygdala activity (Fig. 1c) recorded during the viewing of IAPS images, to evaluate whether the model’s internal workings share similarities with actual brain processes.

### Model-predicted valence and arousal versus normative human ratings

Valence and arousal ratings represent the semantic categorization of affective images by human participants. We compared the model’s predicted valence and arousal against the human ratings of the 1,182 IAPS images (target ratings) that were excluded from the training data. Correlations between predicted and human-rated valence and arousal from these IAPS images are shown in the scatter plots in Fig. 2a. Comparing these correlations before and after training demonstrates the model’s capacity to learn emotional significance from visual features. Prior to training, the amygdala module was randomly initialized using Kaiming initialization, resulting in low or even negative correlation values between predicted and human-rated emotion ratings (valence correlation: –0.2139; arousal correlation: –0.1199). These low correlations are expected, as the model had not yet learned how to assess the emotional significance of the visual input. After training, the valence correlation and the arousal correlation increased markedly to 0.8881 and 0.7003 respectively, demonstrating that the VCA model, in particular the amygdala module, can learn and accurately assess the emotional significance of visual input. The predicted valence range of 1 to 8 spans almost the entire spectrum of possible values between 1 and 9, whereas the predicted arousal range of 3 to 7 shows that the model is less capable of predicting extreme values of arousal.

**Fig. 2.**
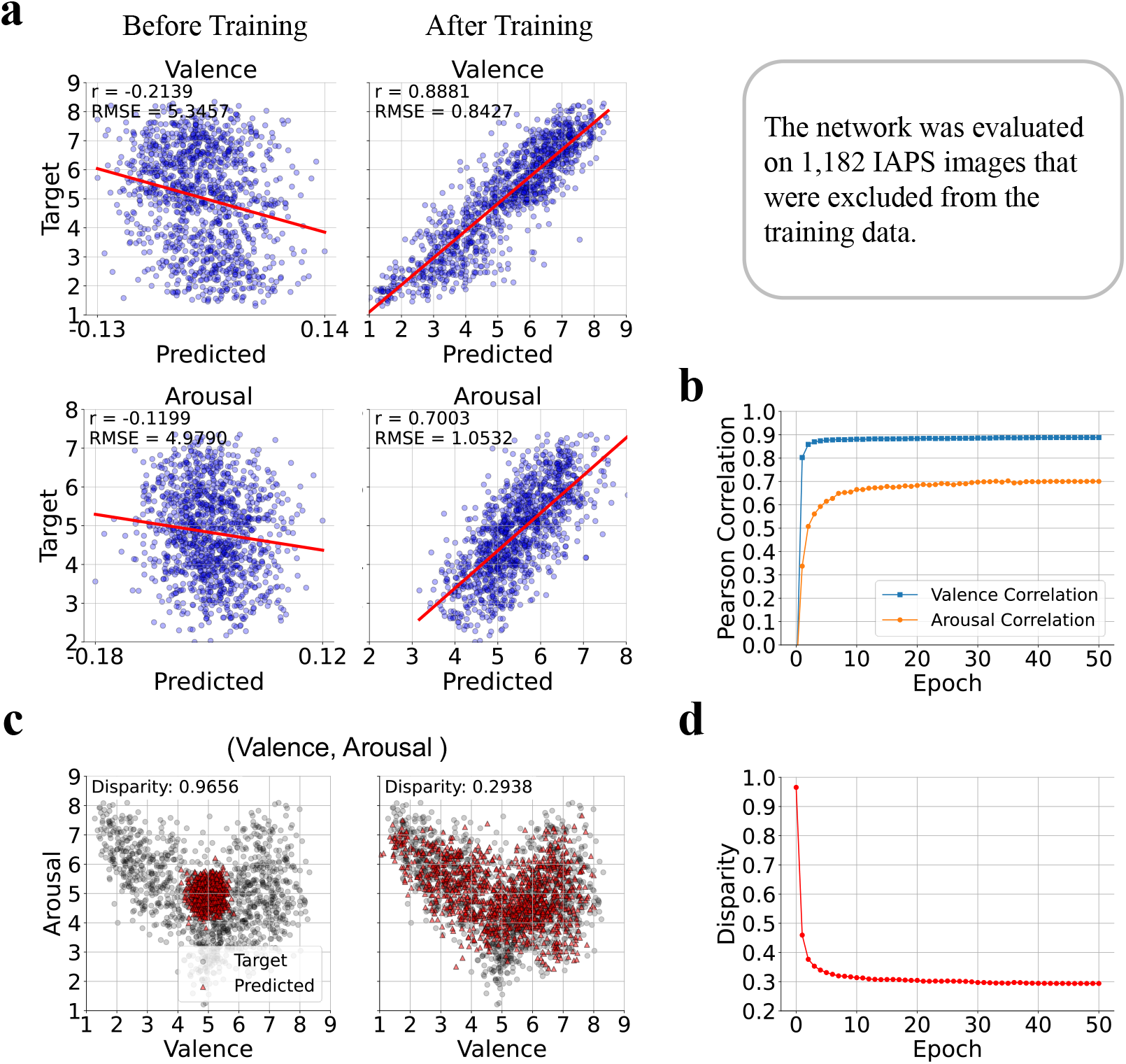
Valence and arousal prediction. **a**, Model predicted valence (top) and arousal (bottom) against target (measured) valence and arousal before (left) and after (right) training. The red line represents the regression fit, and Pearson correlation (r) and RMSE values measure accuracy of prediction. **b**, Pearson correlation of valence and arousal predictions across training epochs. The valence correlation (orange) increases rapidly and stabilizes, while arousal correlation (blue) improves more gradually. **c**, Visualization of model predicted joint valence and arousal representation versus target joint representation before and after training. Procrustes disparity measures the joint accuracy of valence and arousal predictions, with smaller Procrustes disparity value indicating better agreement between predicted valence/arousal and target valence/arousal. Target values (black) and predicted values (red) are shown in the valence-arousal space, with Procrustes disparity value decreasing from 0.9656 to 0.2938 after training, a 70% decrease. **d**, Procrustes disparity reduction across training epochs.

To examine how learning evolves over training, we plotted the correlations between the predicted and human-rated valence and arousal across training epochs (Fig. 2b). The valence correlation increased rapidly, reaching its peak after just a few epochs, while the arousal correlation peaked more slowly, in about ten epochs. This pattern suggests that the model efficiently learns emotional features from visual inputs.

Valence and arousal are not independent variables. They exhibit a V-shaped relationship, reflecting that both high (pleasant) and low (unpleasant) valence images are associated with higher arousal, while mid-range valence corresponds to lower arousal. To assess the model’s joint prediction of valence and arousal, we plotted the Procrustes disparity between the predicted and human ratings before and after training (Fig. 2c). Procrustes disparity quantifies the deviation between two distributions in the two-dimensional valence-arousal space, with smaller values indicating a stronger agreement between the predicted and the actual valence and arousal data. Before training, the model’s predictions clustered near the center of this space, with high disparity (0.9656) between the predicted and the actual data. After training, the disparity dropped to 0.2938, and in particular, the model was able to capture the V-shaped relationship between arousal and valence, consistent with prior psychological studies (Kuppens et al. 2013; Yik et al. 2023). The evolution of joint prediction disparity across training epochs (Fig. 2d) showed that initially high, the Procrustes disparity decreases rapidly and then stabilizes as training progressed.

### Representational alignments between VCA amygdala module and the human amygdala

We utilized representational dissimilarity analysis to quantify and compare the alignment of internal activity between the VCA amygdala module and the human amygdala. The internal activity of the VCA amygdala module was represented by the combined outputs of all layers, while the internal activity of the human amygdala was captured using fMRI data (see Fig.3a). Our hypothesis was that affective representations within the amygdala module would strengthen during training and become more aligned with the human amygdala. To test this, we compared the RDM of our biologically inspired amygdala module with that derived from the human amygdala. Figure 3b displays the Spearman correlation between the two RDMs as blue bars across training epochs. The correlation steadily increased from approximately 0.0 at epoch 0 to 0.12 at epoch 50, indicating a progressively increased alignment between the VCA module representation of IAPS images and the human amygdala representation of the same set of images. This result underscores the model’s capability to capture key aspects of human emotional processing through learning, lending support to its brain-inspired design.

**Fig. 3.**
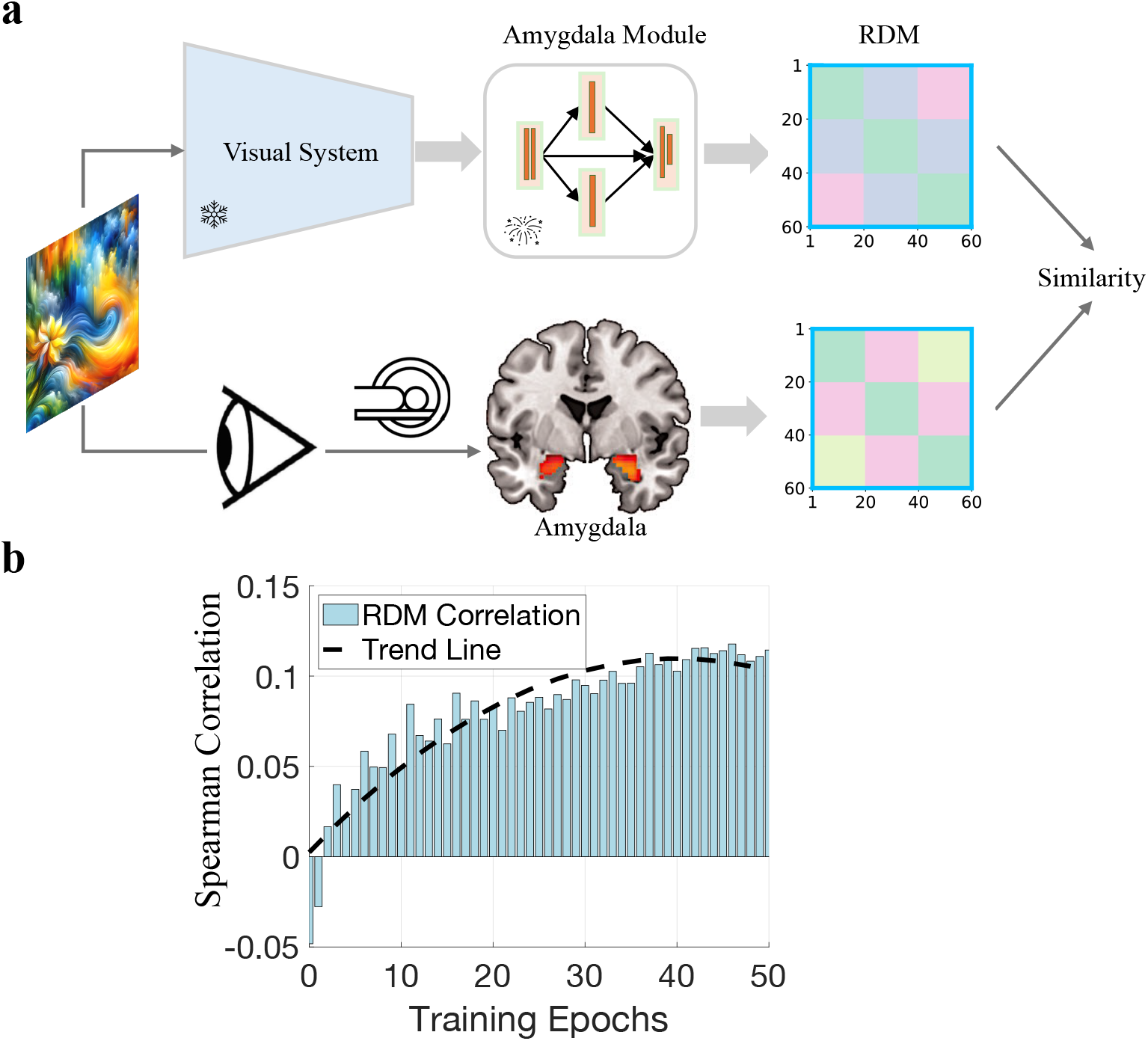
Representational alignment between the VCA amygdala module and human amygdala fMRI activity. **a**, Schematic of the analysis pipeline. Responses in the VCA amygdala module to IAPS images were used to generate the representational dissimilarity matrix (RDM). fMRI responses in the human amygdala to the same images was used to construct fMRI RDM. The similarity between the model and neural RDMs is computed to evaluate representational alignment. **b**, The correlation between model and fMRI RDMs increases over training epochs, indicating improved representational alignment between emotional representations in the model amygdala and biological amygdala. The trend line (black dashed) highlights the overall trajectory.

### Effects of different types of visual modules

The visual representations extracted from the input images by the visual module become the input into the amygdala module. Various network architectures, ranging from shallow to deep and employing different computational methods such as convolution or transformer-based approaches, have been used to model the visual system. How these different architectures impact the emotional assessment have not been systematically studied. Moreover, training methods and training datasets influence the quality and diversity of these representations, which can impact the affective interpretations within the amygdala module. By varying the visual system while keeping the amygdala module constant, with its fixed self-attention and anatomical constraints, we can assess how network architecture and training regimes impact valence and arousal predictions (see Fig. 4a). We hypothesized that rich visual features from CLIP training and self-attention visual systems would yield superior predictions.

**Fig. 4.**
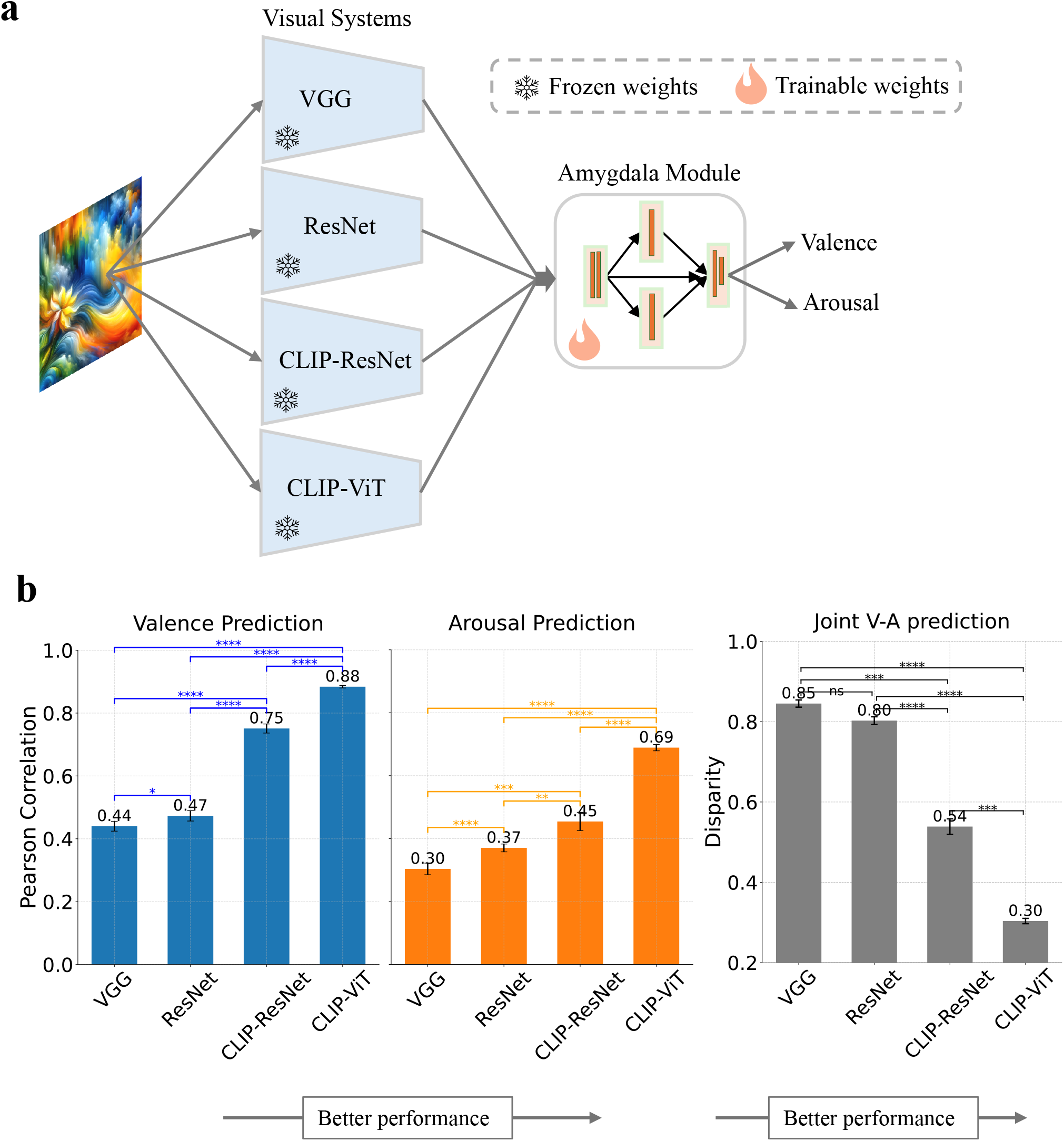
Performance comparison of different visual systems in predicting valence and arousal. **a**, Schematic representation of the experimental setup, where four different visual systems (VGG, ResNet, CLIP-ResNet, and CLIP-ViT) are used to process input images, and their outputs are sent to the amygdala module used above to predict valence and arousal. **b**, Left: Pearson correlation between predicted and actual valence (blue) and arousal (orange) for each visual system after training. Higher values indicate better prediction performance. Right: Procrustes disparity in prediction performance across valence and arousal, with lower values indicating better prediction performance. CLIP-ViT, which is the visual module used in Figures 2 and 3, achieves highest correlation and lowest Procrustes disparity, thereby demonstrating the best performance among the four models. Pairwise comparisons between adjacent models were performed for both results using independent t-tests with Bonferroni correction; Cohen’s d >0.8.

The networks considered include VGG (Sengupta et al. 2019), ResNet (He et al. 2016), CLIP-ResNet, and CLIP-ViT (Radford et al. 2021), with CLIP-VIT being the visual system used the foregoing analysis. VGG and ResNet were pre-trained on ImageNet, a pure image dataset, while CLIP-ResNet and CLIP-ViT were pre-trained on millions of text-image pairs using CLIP. Comparing VGG with ResNet can reveal the impact of shallow versus deep architectures, whereas contrasting CLIP-ResNet with CLIP-ViT allows us to assess the difference between convolution-based and transformer-based self-attention structures. Finally, comparing ResNet with CLIP-ResNet highlights the benefits of large-scale, context-rich pre-training.

Fig. 4b shows that after training, CLIP-ViT outperforms the others in both valence and arousal predictions. Specifically, the arousal correlations are 0.30 for VGG, 0.37 for ResNet, 0.45 for CLIP-ResNet, and 0.69 for CLIP-ViT. Correlations for valence predictions are higher than for arousal predictions: 0.44 for VGG, 0.47 for ResNet, 0.75 for CLIP-ResNet, and 0.88 for CLIP-ViT. These results indicate that deeper networks offer advantages over shallower ones. Moreover, the improvement from ResNet to CLIP-ResNet (0.47 to 0.75 for valence) underscores the benefit of large-scale, context-rich pre-training, even though the gain in arousal prediction is more modest. Comparisons between CLIP-ResNet and CLIP-ViT suggest that transformer-based architectures with self-attention mechanisms excel at capturing semantics-rich visual representations from affective images. To evaluate joint predictions of valence and arousal, we plotted the Procrustes disparity (Fig. 4b right). The results confirm that CLIP-ViT achieves the best alignment with human normative ratings, followed by CLIP-ResNet, ResNet, and VGG.

Our findings demonstrate that varying the visual system significantly influences affective learning. Rich semantic representations, especially those derived from transformer-based architectures and large-scale CLIP training, are critical for accurate valence and arousal predictions, underscoring the importance of network design and pre-training in modeling nuanced emotional cues.

### Effects of different types of amygdala modules

The influence of different structural configurations and computational mechanisms within the amygdala module on valence and arousal prediction is considered next. CLIP-ViT was taken as the visual module and paired with four different types of amygdala modules (see Fig. 5a): Model I – No Anatomical Constraints, No Self-Attention: This design uses four serial fully connected layers without anatomical constraints or self-attention. It serves as a baseline for affective learning. Model II – No Anatomical Constraints, With Self-Attention: This design employs a series of transformer blocks with self-attention but omits anatomical constraints. It tests the isolated impact of self-attention on affective learning. Model III – With Anatomical Constraints, No Self-Attention: In this configuration, the module follows anatomical constraints in its layer connections while relying on traditional fully connected layers. This design assesses the effect of anatomical constraints alone. Model IV – With Anatomical Constraints, With Self-Attention: This design integrates both anatomical constraints and self-attention within transformer blocks to capture the combined benefits of structural fidelity and advanced computational processing. Here by anatomical constraint we mean that the information flows through the amygdala module along two parallel pathways and in a stepwise order (from lateral to basal to central), rather than through unrestricted, all-to-all connections, mirroring how signals flow through various amygdala nuclei. Model IV was the model used for the amygdala module in earlier investigations.

**Fig. 5.**
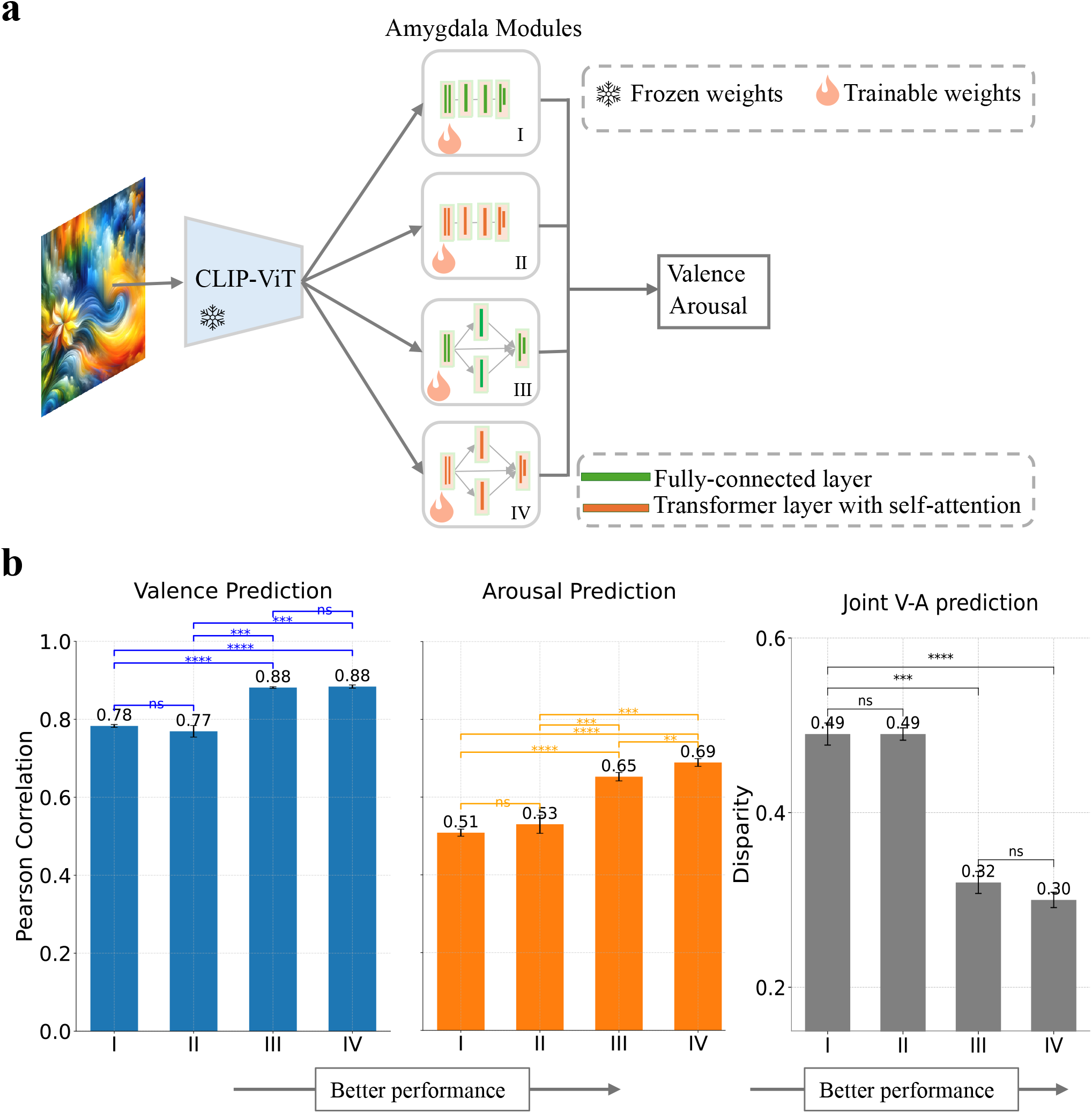
Performance comparison of different amygdala modules in predicting valence and arousal. **a**, Schematic representation of the experimental setup. Four different amygdala modules are: (I) four fully-connected layer blocks connected in a serial manner; (II) four transformer blocks with self-attention connected in a serial manner; (III) four fully-connected layer blocks connected in the anatomical constrained manner; (IV) four transformer blocks with self-attention connected in the anatomical constrained manner. Each module receives visual representations from the CLIP-ViT visual module and is trained to predict valence and arousal. **b**, Left: Pearson correlation between predicted and actual valence (blue) and arousal (orange) for each amygdala model. Higher values indicate better prediction performance. Right: Procrustes disparity in prediction performance across valence and arousal, with lower values indicating better prediction performance. Model IV, which is the model used in the foregoing analysis and which incorporates anatomical constrain and self-attention, achieves the highest prediction performance. Pairwise comparisons between adjacent models were performed for both results using independent t-tests with Bonferroni correction; Cohen’s d >0.8.

Fig. 5b presents the prediction results across all IAPS images for each model variant. Models I and II, which lack anatomical constraints, yield the lowest performance with Pearson correlations of 0.78 and 0.77 for valence, and 0.52 and 0.53 for arousal, respectively. In contrast, Models III and IV, which incorporate anatomical constraints, significantly outperform the others, achieving a valence correlation of 0.88 in both models and arousal correlations of 0.65 and 0.69. These findings underscore the critical role of anatomical constraints in affective learning. Moreover, comparing Models II and IV indicates that self-attention enhances arousal prediction, improving alignment with human normative ratings, while its effect on valence prediction is minimal.

### Emergence of emotion selectivity in the amygdala module

To better understand how learning changed the response properties of the amygdala neurons to emotional input, we tracked the responses of all artificial neurons in the amygdala module to 1,182 IAPS images before and after learning. Each image was assigned to one of nine valence-arousal bins on the valence-arousal plane (Fig. 6a). To quantify whether a neuron is tuned to a particular combination of valence and arousal, we computed an eight-dimensional **d′** (d-prime) profile for each neuron, excluding bins with fewer than ten images. A neuron was labeled emotion selective if its d′ exceeded 0.65, meaning its response to images in the preferred bin was roughly twice that of other bins.

**Fig. 6.**
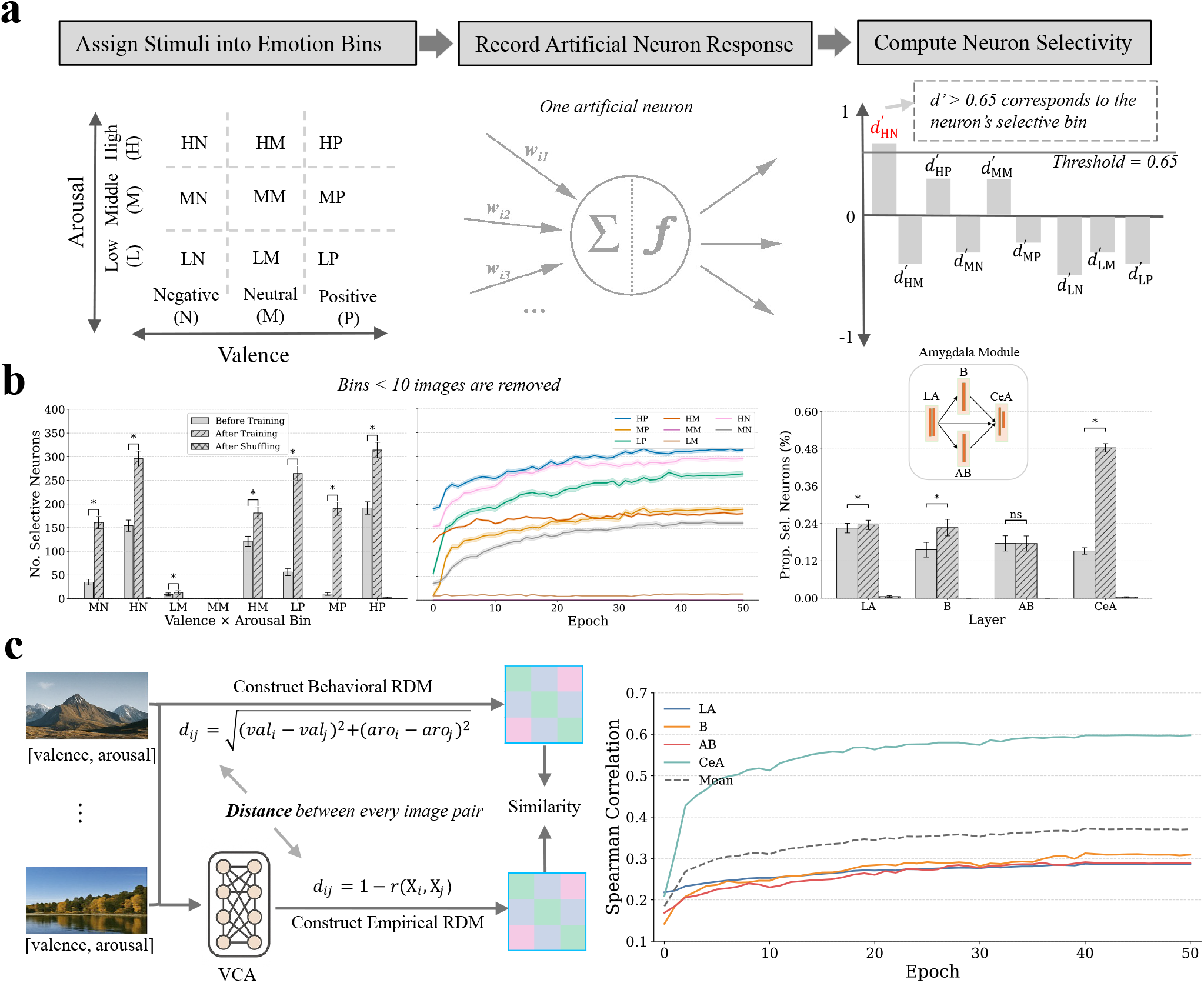
Emergence of emotion selectivity and population-level emotion representational in the amygdala module. **a**, Single-neuron selectivity analysis pipeline. Stimuli are first assigned to one of nine affective bins defined by valence (negative, middle, positive) and arousal (low, middle, high). Artificial neurons’ responses to each stimulus are recorded, and bin-wise selectivity is computed using d-prime (d′). A neuron is considered selective for a bin if its d′ exceeds 0.65. **b**, Training-dependent increase in bin-selective neurons. Left: Number of emotion selective neurons per emotion bin, before and after training and label shuffling. Middle: Proportion of selective neurons by amygdala layer (LA, B, AB, and CeA). Significant increases in emotion selectivity emerge after training (asterisks indicate *p* <0.05). Right: Amygdala layers show different selectivity profiles, with CeA exhibiting the strongest selectivity. **c**, Population-level representational similarity analysis (RSA). Left: Schematic of representational similarity analysis. Behavioral RDMs are constructed from pairwise Euclidean distances in valence-arousal space with human ratings. Empirical RDMs are derived from neural responses in the amygdala module, where each response vector (X_*i*_) represents the activation pattern across all neurons in a given amygdala layer in response to stimulus *i*. Correlation distances between these vectors are used to construct the empirical RDM. Spearman correlation between the empirical RDM and the behavioral RDM quantify the alignment of learned representations with affective structure. Right: Correlation scores increase over training epochs, especially in CeA. Together, training leads to the emergence of emotion-selective neurons (a–b) and also to increasingly structured population representations (c), supporting the amygdala module’s ability to encode affective information at both the single unit and population levels.

Fig. 6b left shows the number of emotion selective neurons in each bin before training, after training, and under label-shuffling control. Nearly all eight valid bins showed a significant post-training increase in emotion selective neurons (asterisks, p < 0.05), while shuffled controls remained near zero. For example, the number of neurons selective for High Arousal-Positive Valence (HP) bin rose from fewer than 200 to over 300. Similarly, neurons selective for High Arousal-Negative Valence (HN) increased from 150 to nearly 290, for Low Arousal-Negative Valence (LN) from about 50 to 270, and for Middle Arousal-Positive Valence (MP) from fewer than 10 to almost 200, which is a 20-fold increase. These results confirm that emotional selectivity emerged through learning, which did not occur by chance.

Over training epochs, as shown in Fig. 6b middle, few neurons were emotion selective at epoch 1, and the number rose steadily, especially for HP, HN, LP, and MP bins, as learning progressed. Bins such as Middle Arousal-Negative Valence (MN) and High Arousal-Middle Valence (HM) showed moderate gains; Low Arousal-Middle Valence (LM) increased only slightly. No neurons became selective for neutral stimuli. These trends suggest that the model prioritizes emotionally intense stimuli, particularly those high in arousal, while discounting the importance of neutral input. This progression parallels the development of emotional tuning in the brain through experience (Leppänen and Nelson 2009; Casey et al. 2019).

We next examined how emotion selectivity is distributed across layers within the amygdala module (Fig. 6b right). We calculated the proportion of selective neurons in each layer before and after training, and under label-shuffling control. Before training, selectivity was uniformly low across layers (<0.24), with the CeA layer at around 0.13. After training, selectivity increased significantly in B and CeA, with CeA showing the largest jump, to nearly 0.50, significantly higher than all other layers (p < 0.05). The B layer increased from 0.15 to 0.23. LA showed a modest increase (from 0.15 to 0.18), while AB remained unchanged. These results suggest that the development of emotional selectivity is not evenly distributed, but rather becomes progressively refined across the hierarchy, culminating in CeA. This mirrors the anatomical organization of the biological amygdala, where deeper nuclei exhibit more specialized emotional responses (Namburi et al. 2015; Janak and Tye 2015)

Having examined the development of emotional selectivity at the single neuron level, we now examine how well population-level responses in the amygdala model capture affective structure. To this end we performed representational similarity analysis (RSA) (Kriegeskorte et al. 2008). We first constructed a behavioral representational dissimilarity matrix (RDM) by using continuous human valence and arousal ratings. Each stimulus was represented as a two-dimensional vector and pairwise dissimilarity between stimuli was quantified as the Euclidean distance in this affective space. This approach provides a continuous, graded model of emotional similarity. In parallel, we computed empirical RDMs based on correlation distances between neural population responses in each sublayer of the amygdala module (Fig. 6c, left). The Spearman correlation between behavioral and empirical RDMs provided an RSA score, quantifying how closely the model’s internal representational geometry aligns with the behavioral RDM. As shown in the right panel of Fig. 6c, RSA scores increased steadily over training epochs in all four sublayers, LA, B, AB, and CeA, indicating that learning gradually aligns population-level neural representations with the affective structure of human judgments. The most substantial gains occurred in the CeA, which ultimately achieved the highest RSA values (∼0.6), suggesting that this sublayer develops the most behaviorally aligned representations. In contrast, early sublayers (LA, B, AB) showed modest increases and plateaued at lower RSA scores. This layer-wise gradient of alignment implies a functional hierarchy within the amygdala model, whereby the deeper layers encode increasingly more abstract and behaviorally and semantically relevant features.

## Materials and Methods

### Affective datasets

In this study we used affective scene images from four datasets: VideoFrame, IAPS, NAPS, and Kim. The VideoFrame dataset comprises 2,184 images, curated by selecting one frame every ten frames from published affective videos (Cowen and Keltner 2017). The decimation method helped reduce redundancies in the selected images. IAPS is a well-known dataset widely used in affective neuroscience (Davidson 1998; Gusnard et al. 2001; Yoo et al. 2007; Bradley and Lang 2007) (Davidson 1998; Gusnard et al. 2001; Yoo et al. 2007); it contains 1,182 images portraying a broad range of emotions. The NAPS dataset includes 1,356 images and is also employed frequently in affective computing and emotion research (Marchewka et al. 2014; Lehmann et al. 2016; Koush et al. 2017; Behnke et al. 2020). The Kim dataset, consisting of 10,766 images, was introduced in a deep neural network study focused on emotion representation in the human brain (Kim et al. 2018). All images in these datasets have associated normative ratings of valence and arousal. For our study, we combined VideoFrame, NAPS, and Kim datasets to train the VCA model to recognize the valence and arousal of each image, and evaluated the model performance on the IAPS dataset.

### The VCA model

#### The visual module

The proposed visual cortex-amygdala (VCA) model consists of a visual module and an amygdala module. The visual module is the CLIP-trained ViT pretrained on image-text pairs. Three additional deep neural networks, pretrained either on ImageNet or on image-text pairs, were also tested as possible candidates for the visual module to examine the influence of visual encoding on emotional inference. These models represent two distinct approaches to computational operations for extracting and encoding visual semantics from stimuli: convolution-based architectures (e.g., VGG16 and ResNet50) and transformer-based architectures (e.g., ViT).

The rationale for selecting different types of architectures and pretraining methods for the visual system is twofold. First, the quality of visual semantics encoded by the visual system is crucial for effective visual emotion perception. Without a rich and diverse representation of visual semantics, the amygdala module cannot accurately interpret and transform these visual features into affective meanings. Therefore, it is necessary to investigate which architectures and training strategies produce superior visual semantics, ultimately leading to more accurate affective decoding by the amygdala. The exploration of various visual system architectures and their interactions with the amygdala module under different conditions provides a comprehensive understanding of how visual information is transformed into affective representations.

#### The amygdala module

The architecture of the proposed amygdala module is inspired by the gross anatomy of the human amygdala. The model consists of four distinct layers, each representing a corresponding nucleus in the amygdala: the lateral nucleus (LA), accessory basal nucleus (AB), basolateral nucleus (B), and central amygdala (CeA). These layers are connected to mirror the known connection observed in the biological amygdala (Pitkänen et al. 1997; Sah et al. 2003). A key feature of the amygdala module is that signals travel along parallel pathways rather than a single straight line. The LA receives sensory inputs from cortical and subcortical sources and then sends this information through both the basal and accessory basal nuclei before it reaches CeA. In the brain, this parallel organization allows different kinds of information (e.g., sensory signals, contextual cues, and modulatory feedback) to be integrated across nuclei. This organization is thought to make affective processing more flexible and reliable by allowing different groups of neurons to be engaged depending on the situation (Janak and Tye 2015; Gothard 2020). Recent work further suggests that this architecture supports multidimensional and context-dependent representations, enabling the amygdala to guide nuanced emotional evaluations and social decisions (Gothard 2020; Putnam and Gothard 2019).

In addition to these anatomical considerations, we introduced a self-attention mechanism within the amygdala module. Specifically, we utilized transformer blocks to model each layer, wherein each transformer block incorporates self-attention mechanisms. The rationale behind the introduction of the self-attention mechanism stems from the presence of astrocytes in the biological amygdala, which have been hypothesized in some studies to perform computations analogous to self-attention.

Similar to the consideration of different models for the visual system we also considered several other models of the amygdala module. In Model I, the amygdala module comprises four fully connected layers arranged sequentially, which does not incorporate anatomical constraints and self-attention. In Model II, the module is composed of four transformer blocks employing multi-head self-attention, also connected in series; this model incorporates self-attention but no anatomical constraints. Model III uses fully connected layers where the interconnections are guided by the gross anatomy of the biological amygdala but does not include self-attention. Finally, Model IV combines transformer blocks with self-attention and a biologically constrained connectivity architecture, which is described in more detail in the previous paragraph.

#### Mathematical formulation

The mathematical formulation of the proposed VCA model shown in Figure 1a is as follows. Let *X* be a visual image input. The function *VisualSystem*(*x*) denotes a pretrained deep neural network for visual processing (e.g., CLIP-trained ViT), producing an output *y*_*visual*_ as the representation of the input image:

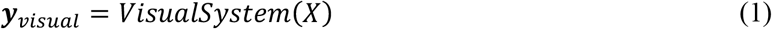

This output is passed to the LA layer of the amygdala module, where self-attention is computed and followed by a linear transformation with learnable weights ***W***_*LA*_ and ***b***_*LA*_:

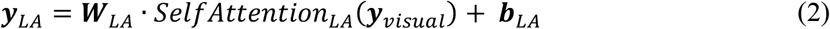

The processed representation *y*_*LA*_ then feeds into two parallel branches: the AB layer and the B layer. Each applies its own self-attention mechanism and linear transformation:

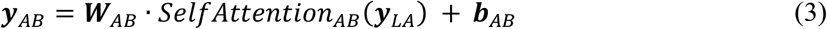

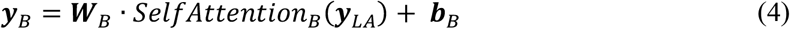

Next, the outputs from LA, AB, and B layers are concatenated and processed by the CE layer, which performs self-attention followed by a linear transformation:

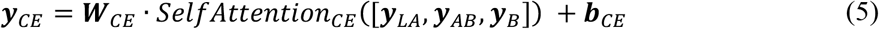

Finally, the resulting vector *y*_*CE*_ is passed through a linear regression layer to generate the final output prediction *y*:

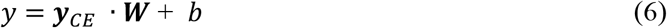

Here, SelfAttention refers to a self-attention operation applied within each layer in the amygdala module., operating on the input it receives from the previous stage. Each weight matrix ***W***_*∗*_ and bias ***b***_*∗*_ is learned during training and specific to its corresponding layer.

More details of self-attention are given below.

For a given input *X*∈ ℝ^*n*×*d*^, where *n* is the number of tokens and *d* is the feature dimension, the self-attention mechanism performs three core computations. First, the input is projected into three separate spaces using learnable weight matrices to produce the query ***Q***, key ***K***, and value ***V*** matrices using learnable weight matrices:

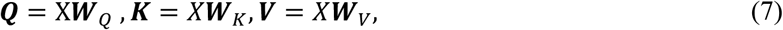

Here, 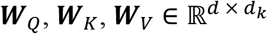 are trainable projection matrices, and *d*_*k*_ is the dimensionality of the key and query vectors.

The self-attention output is computed using the scaled dot-product attention formula:

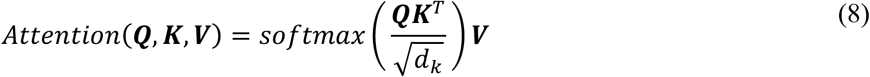

This formulation enables each token to attend to all other tokens in the sequence. The scaling factor 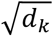 stabilizes the gradients by preventing the dot-product values from becoming excessively large, which would otherwise saturate the softmax function.

Second, it is the feedforward neural network (FFN) that applies two linear transformations with a non-linear activation (ReLU) in between. It can be formulated as

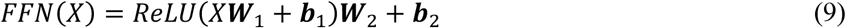

The third computation is layer normalization and residual connections and can be formulated as

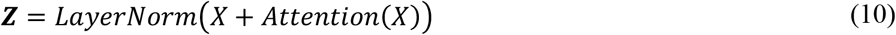

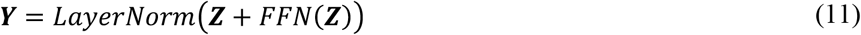

Here, *LayerNorm* normalizes the inputs across the features for each data sample. Given an input vector ***h*** ∈ ℝ^*d*^ for a single data sample with d dimensions of features, it computes the normalized output as below:

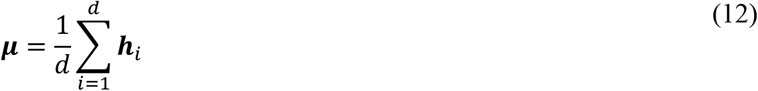

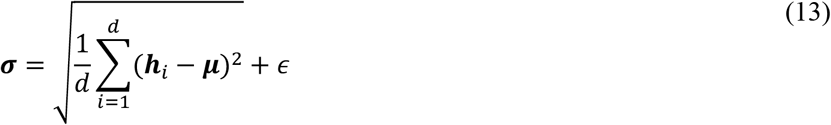

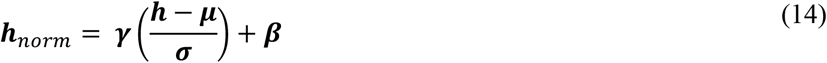

where *μ* is the mean of the features for the data sample; σ is the standard deviation of the features, with ϵ = 10^−5^ added for numerical stability; **γ, β** ∈ ℝ^*d*^ are learnable parameters that scale and shift the normalized output; ***h***_*norm*_ ∈ ℝ^*d*^ are normalized output features.

The development of DNNs includes defining an objective function, image preprocessing, training, and testing. We use mean squared error (MSE) as the objective function, which can be formulated as:

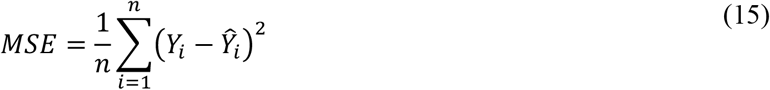

where *n* is the number of images used for training; *Y*_*i*_is the true valence value and Ŷ_*i*_is the predicted valence for the *i*-th image. ***Y*** ∈ ℝ^*n*^ and Ŷ∈ ℝ^*n*^ are the vectors of ground-truth and predicted values, respectively. Essentially, it is a regression problem using MSE as a loss function to calculate the difference between DNNs’ predicted valence/arousal and the true valence/arousal rated by human subjects. The difference is the prediction error that is used for updating the weights of DNNs through back-propagation.

#### Preprocessing and Training

We applied a series of image preprocessing steps prior to training the DNN. In particular, each image was resized to 224 x 224 pixels, rotated by 20 degrees with a probability of 0.5, and randomly flipped horizontally with a probability of 0.5 The rotation and flipping were applied only during the training, while the resizing step were applied during both training and testing. These preprocessing steps help reduce risk of overfitting in DNN training. For model training, only the amygdala modules were trainable, while the visual system modules had fixed weights. We trained the DNNs using a batch size of 64, an initial learning rate of 1e-4, and stochastic gradient descent (SGD) for optimization over 50 epochs. A weight decay of 1e-4 was applied for regularization. The learning rate was reduced by a factor of 0.1 every 40 epochs to facilitate convergence.

### fMRI data

#### fMRI Experimental Paradigm

A previously published dataset was used here (Bo et al. 2021). Twenty-six healthy volunteers with normal or corrected-to-normal vision gave written informed consent and participated in the study. The fMRI study protocol was approved by the University of Florida Institutional Review Board. Prior to the MRI scan, two participants withdrew from the study. Data from an additional four participants were excluded due to excessive head motion during scanning. As a result, data from the remaining twenty participants were included in the final analysis (mean age: 20.4±3.1 years; 10 male, 10 female).

During the experiment, participants were instructed to passively view a series of affective pictures from IAPS while simultaneous EEG-fMRI data were recorded (EEG data is not analyzed in the current study). Each IAPS picture was displayed on a MR-compatible monitor for 3s, which was followed by a variable interstimulus interval (ISI) of either 2800 ms or 4300 ms. The use of variable ISIs helps reduce temporal predictability and allows for better estimation of the hemodynamic response in event-related fMRI designs. Each session comprised 60 trials corresponding to 60 different pictures broadly divided into three categories: pleasant, neutral and unpleasant. There were five sessions and the order of the same 60 pictures was randomized from session to session. Pictures were presented on the monitor that was placed outside the scanner and viewed via a reflective mirror. Participants were instructed to maintain fixation on the center of the screen during the whole session.

#### Acquisition and Preprocessing

Functional MRI data were collected using a 3T Philips Achieva scanner (Philips Medical Systems). The scanner parameters were as follows: echo time (TE) = 30 ms, repetition time (TR) = 1.98 s, flip angle= 80°, slice number = 36, field of view = 224 mm, voxel size = 3.5 × 3.5 × 3.5 mm^3^, and matrix size = 64 × 64. Slices were acquired in ascending order and oriented parallel to the anterior-posterior commissure (AC-PC) plane. A high-resolution T1-weighted structural image was also collected. EEG data were simultaneously acquired but are not analyzed in the present study.

Preprocessing of the fMRI data was conducted using Statistical Parametric Mapping (SPM, http://www.fil.ion.ucl.ac.uk/spm/). The first five volumes of each session were discarded to eliminate artifacts caused by the initial transient instability of scanner. Slice timing correction was applied using interpolation to account for differences in acquisition times across slices. Realignment is then performed to correct for head motion across six degrees of freedom (three translations and three rotations). The images were subsequently normalized and registered to the Montreal Neurological Institute (MNI) template, and resampled to a spatial resolution of 3 × 3 × 3 mm^3^. A Gaussian spatial smoothing filter with a full width at half maximum (FWHM) of 8 mm was applied. Finally, a high-pass temporal filter at 1/128Hz was used to remove low frequency temporal drifts.

#### Single-trial Beta-series Estimation

Trial-by-trial picture-evoked responses were estimated using the beta-series method (Mumford et al. 2012). For each trial, a general linear model (GLM) was constructed in which the target trial was modeled with a single regressor, while all other trials were grouped into a separate regressor. Additionally, six motion regressors were included to account for movement-related artifacts. This procedure was repeated for each trial, resulting in a single-trial beta estimate (beta-series) for each stimulus presentation. To improve signal stability, beta-series estimates corresponding to the same picture stimulus presented in different sessions were averaged, yielding sixty unique picture representations per participant for subsequent representational similarity analysis (RSA).

### Representational similarity analysis

We employed RSA to investigate the functional associations between the human amygdala and the amygdala module in the VCA model. The same sixty IAPS images used in the fMRI experiment were also used for obtaining responses in DNN models. To compute the representational dissimilarity matrices (RDMs) from the fMRI data, we first aligned the fMRI data from the amygdala region with the corresponding image indices. The data underwent preprocessing and normalization via z-scoring across voxels for each image and subject. The normalized activation patterns were then averaged across all subjects to generate a mean neural representation for each image. Following this, pairwise Spearman correlations were calculated between the averaged activation vectors for each image. The resulting similarity values were transformed into dissimilarity scores by subtracting the correlation values from one (1 - correlation). More specifically, given the number of voxels *V* = 291 in the amygdala region, the number of images *I* = 60, and the number of subjects *S* = 20, the RDM was constructed according to the following procedure.

For each subject *s* and image *i*, we normalize the voxel activation ***v***_*i,s*_∈ ℝ^*V*^ as:

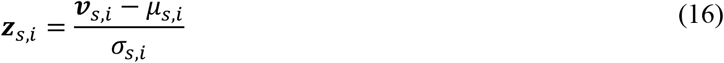

where *μ*_*s,i*_and σ_*s,i*_ are the mean and standard deviation of the non-NaN voxel activations for subject *s* and image *i*. We then compute the average normalized activation across all subjects for each voxel and image:

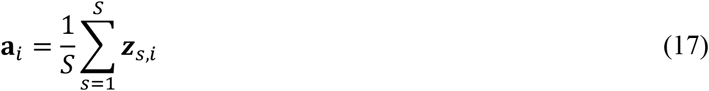

where **a**_*i*_∈ ℝ^*V*^ is the averaged activation vector for image *i*. For each pair of images (*i, j*), we compute the Spearman correlation:

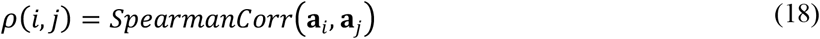

and derive the RDM

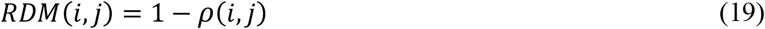

To calculate the RDMs for DNN data, we employed a systematic process consisting of feature extraction, dimensionality reduction, normalization, and dissimilarity assessment. At each training epoch, we extracted feature vectors from multiple sublayers within the VCA model’s amygdala-mimetic module. Specifically, for each model checkpoint corresponding to a particular epoch, we obtained feature representations from four key sublayers: LA, AB, B, and CeA. For the IAPS test images (*N* = 60), each feature matrix *F*_*i*_(where *i* ∈ {LA, AB, B, CeA}) had dimensions *N* × *F*_*i*_, where *F*_*i*_denoting the number of features in the *i*-th layer. Due to the high dimensionality of the combined feature space, we applied principal component analysis (PCA) to reduce the dimensionality while preserving the majority of variance. Specifically, feature vectors were projected onto a *P* = 1,000 principal component space. To ensure comparability across features, z-score normalization was performed on each layer’s reduced feature matrix, standardizing feature dimensions to have zero mean and unit variance. Finally, we quantify the pairwise dissimilarity between all image pairs using Spearman’s rank correlation distance (1-Spearman’s *ρ*) as the metric, chosen for its robustness to non-linear relationships and its capacity to capture monotonic relationships between feature vectors. The resulting RDMs reflect the evolving internal representations within each amygdala sublayer across training.

Finally, we quantified the similarity between DNN-derived representations and a reference amygdala-based RDM across training epochs. Spearman’s rank correlation was used to compare the dissimilarity structures of the two RDMs. This analysis provides insights into how the DNN’s internal representations progressively evolve to align with human neural patterns in the amygdala.

### Emotional selectivity and RSA in the amygdala module

To assess neural selectivity for emotion-related dimensions, we analyzed the responses of each neuron within the amygdala module to 1,182 affective images from the IAPS database. Each image was labeled with valence and arousal scores and passed through the model to extract neural responses from four sublayers of the amygdala module (LA, BL, BA, and CeA). This allowed us to characterize how individual units across different anatomical subcomponents encode affective dimensions.

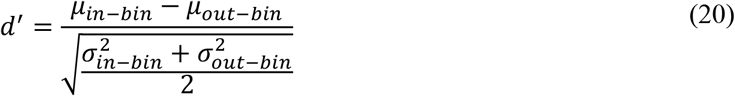

To determine statistical significance, we performed a two-sample *t*-test with unequal variance (Welch’s *t*-test) for each bin-wise comparison. A neuron was considered selective for a given bin if: (1) the difference in responses was statistically significant (*p* < 0.05), and (2) the d′ value exceeded a threshold of 0.65., which has been used in other studies (Aparicio et al. 2016; Lee et al. 2020).

To quantify changes in emotional selectivity, we compared the number of emotional selective neurons across three conditions: Before Training, After Training, and After Shuffling (random permutation of assigned bins to each image). Bootstrap resampling (1,000 iterations) was used to estimate standard errors and assess statistical differences across conditions (Fig. 6b, left). To examine how selectivity evolved throughout training, we tracked the number of selective neurons across training epochs. At each epoch, neurons were evaluated using the same bin-wise d′ and significance criteria described above, and the number of selective neurons was aggregated across layers (Fig. 6b, middle). To determine where in the network these selective neurons emerged, we analyzed the distribution of selectivity across sublayers of the amygdala module. For each layer, we calculated the proportion of neurons that were selective for any bin and compared these proportions across conditions using bootstrap statistics (Fig. 6b, right).

To quantify how well the population activity in each layer of the amygdala module encoded affective structure, we performed RSA between behavioral and empirical RDMs.

#### Behavioral RDM

Constructed from the continuous human behavioral valence and arousal ratings. For each stimulus *i* we form the 2D vector [*val*_*i*_, *aro*_*i*_], and define pairwise dissimilarity as the Euclidean distance in this space:

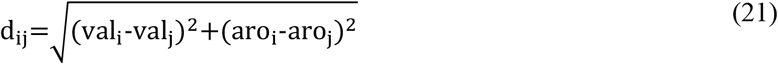

#### Empirical RDMs

For each analysis, stimulus-by-neuron activation patterns were assembled by concatenating units across the selected amygdala layers. Pairwise dissimilarity between stimuli *i* and *j* was computed as correlation distance between their activation vectors using the Eq.18.

For each epoch, we computed the Spearman rank correlation between the upper triangular entries of the behavioral and empirical RDMs, yielding an RSA score for each layer. This procedure was repeated across training epochs, and RSA scores were averaged and visualized with standard error shading to reveal trends in how affective information became embedded in the network over time. To capture an overall summary, we also computed a cross-layer average RSA trajectory (Fig. 6c).

## Discussion

In this study, we developed and evaluated a biologically inspired deep neural network model of visual emotion processing. The model, referred to as the Visual Cortex Amygdala (VCA) model, is composed of two primary modules: a vision transformer module represents the visual system and an amygdala-mimetic module, incorporating anatomical constraints and self-attention blocks, represents the affective module. Trained on three affective image datasets (NAPS, VideoFrame, Kim) and tested on a fourth dataset (IAPS), the model was shown to achieve strong performance at predicting an input image’s valence and arousal, individually as well as jointly. In addition, with learning, the internal representations of the affective inputs in the amygdala module exhibited increased alignment with that of the biological amygdala as assessed by fMRI, underscoring the model’s biological plausibility. Two additional research questions were also considered. First, we examined how variations in the architecture and training paradigm of the visual system influence emotional interpretation, and found that CLIP-ViT, trained on image-text pairs, generated the richest visual representations, which significantly improved prediction of both valence and arousal compared to alternative configurations such as convolutional neural networks trained on ImageNet. Second, we investigated how anatomical and computational constraints within the amygdala module affected model performance, and found that anatomical constraints enhanced the model’s accuracy in predicting both valence and arousal, while attention-based mechanisms further improved arousal prediction. This latter result is particularly noteworthy in light of biological evidence that astrocytes in the amygdala are engaged by arousal-related neuromodulators such as norepinephrine and can shape circuit responses during high-arousal states (e.g., fear) (Bender et al. 2016; Martin-Fernandez et al. 2017; Suthard et al. 2023).

### Visual encoding and emotion assessment

Recognizing the motivational significance of visual input is critical for the survival of an organism. Converging neuroscience evidence indicates that cortical visual encoding of sensory input is essential not only for decoding object information but also for assessing its emotional significance. Anatomical and physiological findings in both primates and human reviewed by (Pessoa and Adolphs 2010) show no strong evidence for the amygdala to perform visual encoding directly. Instead, the amygdala receives its predominant visual input from higher-order visual cortical areas in the anterior temporal lobe, where signals contain identity, category, and high-spatial-frequence details (Kreiman et al. 2000; Gothard et al. 2007; Mormann et al. 2008). These rich cortical representations provide the informational foundation for the amygdala’s modulatory role in guiding attention, perception, and behavior toward emotionally relevant stimuli. Thus, the ability of the amygdala to assign motivational values to input depends critically on the robustness and precision of cortical visual encoding. Supporting this view, (Conwell et al. 2025) systematically evaluated 180 state-of-the-art deep neural networks trained on computer vision tasks (e.g., object recognition), and found that learned visual features, whether from natural images or paired image–text data, can explain a majority of the explainable variance in human ratings of arousal, valence, and beauty. Particularly, vision–language models significantly outperformed pure vision models (mean r^2^ = 0.87 vs. 0.72), underscoring that rich, experience-dependent perceptual representations carry substantial affect-predictive information.

Our findings lend further support to the critical role of visual encoding in shaping emotional interpretation. Richer visual representations, particularly those learned through multimodal data (e.g., image-text pairs), enabled more accurate prediction of emotional responses. These results align with prior studies showing that visual features extracted from deep learning models trained on scene or object recognition tasks can explain substantial variance in human ratings of emotion and aesthetics (Conwell et al. 2025). Moreover, the superior performance of the models trained with vision-language alignment (e.g., CLIP) suggests that semantic grounding enhances affective interpretation, consistent with the well-established notion that valence and arousal reflect semantic categorization of visual scenes (Bradley and Lang 2007; Nummenmaa et al. 2010; Kuppens et al. 2013; Sadeghi et al. 2024). From a computational modeling perspective, pairing images with natural language captions forces the model to align visual representations with the rich, structured semantic space of language. This joint training not only preserves low- and mid-level perceptual structures but also embeds higher-order conceptual dimensions, such as social meaning, symbolic reference, and emotional tone, that humans use when evaluating visual content (Conwell et al. 2025). In computational models, this alignment has been shown to yield robust, semantically grounded features with strong generalization across tasks and domains (Radford et al. 2021; Li et al. 2022), improve invariance to superficial changes and category variability (Yuksekgonul et al. 2023), and enhance sensitivity to abstract and relational cues (Alayrac et al. 2022). More recently, (Doerig et al. 2025) revealed that high-level visual representations in the human brain are aligned with large language models trained exclusively on text, further underlining the synergistic nature of visual and semantic processing.

### Structural and computational constraints on amygdala function

In the brain, function arises from structure (Sporns 2011; Lichtman and Sanes 2008; Bassett and Sporns 2017; Zador 2019). The amygdala, because of its layered subnuclei and extensive connectivity, is uniquely suited for emotion processing (Sah et al. 2003; Janak and Tye 2015). For instance, the basolateral complex assigns affective salience to sensory input (Phelps and LeDoux 2005; Duvarci and Pare 2014), while the central nucleus drives behavioral and autonomic responses (LeDoux 2000; Viviani et al. 2011). These regions coordinate both rapid, unconscious emotional reactions via subcortical pathways and more deliberative evaluations through cortical inputs (Pessoa and Adolphs 2010). Deep neural network models of the amygdala have thus far not systematically incorporated the anatomical constraints (Kragel et al. 2019; Jang and Kragel 2025). Inspired by recent advances demonstrating the power of structure-constrained models (Lappalainen et al. 2024), we introduced structural constraints into the amygdala module, drawing on the gross anatomy of the biological amygdala, particularly the parallel pathways connecting input to the output. In the brain, parallel routes in the amygdala are thought to allow sensory and contextual information to converge across nuclei, potentially supporting flexible, context-dependent affective processing (Pitkänen et al. 1997; Sah et al. 2003; Gothard 2020) In our model, visual inputs pass through layers simulating the lateral, basal, and central nuclei, enabling progressive more abstract representations of affective significance. Our results show that architectural differences within the amygdala-inspired module significantly influence the model’s ability to predict emotional valence and arousal (Fig. 5). Specifically, Model IV, which incorporates both anatomical constrains and transformer-based self-attention mechanisms, consistently outperforms the other variants across all three performance metrics: valence prediction, arousal prediction, and joint valance-arousal prediction, highlighting the functional advantage of biologically grounded design.

While structure defines the routes through which signals can travel, computation governs how those signals are transformed along the way. Inspired by emerging evidence implicating astrocytes in neural computation, we incorporated self-attention mechanisms into the amygdala module of our model. Traditionally viewed as passive support cells, astrocytes are now understood to modulate synaptic activity through processes like calcium signaling and tripartite synapse dynamics, suggesting a role in implementing context-dependent computations (Perea et al. 2014; Khakh and Sofroniew 2015; Yu et al. 2018). Notably, one recent study (Kozachkov et al. 2023) suggests that astrocytes may naturally implement self-attention-like processes, effectively approximating core computational principles underlying Transformer architectures. Our results demonstrate that models integrating anatomical constraints with attention mechanisms achieve the highest accuracy in predicting emotional valence and arousal, as well as superior learning efficiency. These models converge more quickly during training and exhibit superior generalization, echoing predictions from astrocyte-based computational theories. By uniting biologically inspired structural organization with self-attention-based computation, our model offers a principled framework for studying emotion perception. It advances both the functional capabilities of artificial systems and our understanding of the computational motifs underlying emotional processing in the brain.

### Neural mechanisms of emotion representation in the amygdala module

We make several observations concerning how the amygdala, through learning, acquires the ability to assess the emotional significance of visual input. First, at the single neuron level, we observe the emergence of *emotion selectivity*, where neurons develop strong preference for specific combinations of valence and arousal (Fig. 6). Specifically, dividing the valence-arousal plane 9 different bins, the bin-wise tuning emerges strongly after training, with more selective neurons developing for high-arousal negative (HN) and high-arousal positive (HP) bins, indicating that the module prioritizes emotionally salient inputs. This aligns with studies in rodents and primates showing that amygdala neurons integrate multiple affective signals in nonlinear, context-dependent ways, supporting flexible emotional representations (Janak & Tye, 2015; Fadok et al., 2017). Second, the model develops a clear *bias toward high-arousal inputs*, with disproportionately more neurons tuned to high-arousal bins, particularly the High Arousal-Positive Valence (HP) bin and the High Arousal-Negative Valence bin, after training (Fig. 6b, left and middle). This mirrors findings that the biological amygdala is especially sensitive to highly arousing stimuli whose rapid and accurate representation is essential to guide decision-making and action (LeDoux, 2000; Anderson & Phelps, 2001). Neurophysiological work has shown that neurons in the basolateral and intercalated amygdala respond more strongly and persistently to high-arousal cues (Zhang & Li, 2018). Our model similarly reveals that arousal selectivity becomes organized within specific subpopulations, suggesting that arousal forms a dominant axis of affective representation (Gründemann et al., 2023). Third, we find strong evidence for *experience-dependent plasticity*. The number of emotion-selective neurons increases substantially after training (Fig. 6b, right). These results echo findings from biological systems, where emotional learning shapes selectivity through synaptic plasticity and repeated exposure (Kim & Jung, 2006). Fourth, to probe how affective structure is captured at the *population level*, we performed representational similarity analysis between behavioral RDMs (based on distances on the valence-arousal plane) and empirical RDMs (based on activation patterns; Fig. 6c). RSA scores increased over training, especially in the CeA, indicating that representations in deeper layers become increasingly aligned with the underlying affective geometry. This learning-dependent shift reflects the model’s capacity for experience-dependent plasticity, a hallmark of emotional learning in biological systems (McGaugh 2004; Pessoa and Adolphs 2010).

### Limitations and future directions

Despite these contributions, several limitations warrant consideration. First, our study focused on feedforward architectures and did not incorporate feedback and recurrent neural dynamics, limiting the model’s biological realism. Future models may benefit from incorporating recurrent or predictive coding frameworks to better approximate the brain’s dynamic processing of affective stimuli. Second, the present study focuses exclusively on visual scene-based emotion perception, without accounting for other situations such as perceiving emotion from facial expressions, body language, written and spoken language, or auditory cues, each of which plays a central role in naturalistic emotional understanding. Expanding the model to integrate multiple input types, including auditory or linguistic information, could make it more realistic and help explain emotional perception more effectively. Third, emotional perception is inherently subjective. Individuals may experience distinct affective responses to the same stimulus due to differences in prior experience, personality, or context. Our current model does not capture such individual variability. Future work should consider modeling individual differences, perhaps through personalized embeddings to better reflect the heterogeneity of emotional experience. Fourth, although our amygdala-inspired module attempts to incorporate known anatomical features, it necessarily abstracts many biological complexities. These include cellular diversity (e.g., multiple neuron and glial subtypes), neuromodulatory influences, and dynamic plasticity processes. Future work should aim to incorporate these dimensions to further enhance the biological fidelity and functional insight of computational models of emotion.

## Summary

This study introduced the VCA model that can assess the motivational significance of visual input. By combining a CLIP-pretrained vision transformer with an amygdala-inspired module, we demonstrated that emotional valence and arousal can be accurately and robustly predicted from visual scene input, and with learning, the activity patterns in the amygdala module showed increased alignment with fMRI response patterns in the human amygdala. In addition, the neural mechanisms of emotion assessment at both the single neuron and neuronal population level were examined, and the importance of semantics-rich visual representations and the incorporation of anatomical and computational constraints highlighted. Limitations and future directions were also pointed out.

